# The Type VI secretion system *of Stenotrophomonas rhizophila* CFBP13503 limits the transmission of *Xanthomonas campestris* pv*. campestris* 8004 from radish seeds to seedlings

**DOI:** 10.1101/2023.07.21.549874

**Authors:** Tiffany Garin, Chrystelle Brin, Anne Préveaux, Agathe Brault, Martial Briand, Marie Simonin, Matthieu Barret, Laure Journet, Alain Sarniguet

## Abstract

*Stenotrophomonas rhizophila* CFBP13503 is a seed-borne commensal bacterial strain, which is efficiently transmitted to seedlings and can outcompete the phytopathogenic bacteria *Xanthomonas campestris* pv. *campestris* (Xcc8004). The type VI Secretion System (T6SS), an interference contact-dependent mechanism, is a critical component of interbacterial competition. The involvement of the T6SS of *S. rhizophila* CFBP13503 in the inhibition of Xcc8004 growth and seed-to-seedling transmission was assessed. The T6SS cluster of *S. rhizophila* CFBP13503 and nine putative effectors were identified. Deletion of two T6SS structural genes, *hcp* and *tssB*, abolished the competitive advantage of *S. rhizophila* against Xcc8004 in *vitro*. The population sizes of these two bacterial species were monitored in seedlings after inoculation of radish seeds with mixtures of Xcc8004 and either *S. rhizophila* wild type (wt) strain or isogenic *hcp* mutant. A significant decrease in the population size of Xcc8004 was observed during confrontation with the *S. rhizophila* wt in comparison to T6SS- deletion mutants in germinated seeds and seedlings. We found that the T6SS distribution among 835 genomes of the *Stenotrophomona*s genus is scarce. In contrast, in all available *S. rhizophila* genomes, T6SS clusters are widespread and mainly belong to the T6SS group i4. In conclusion, the T6SS of S. *rhizophila* CFBP13503 is involved in the antibiosis against Xcc8004 and reduces seedling transmission of Xcc8004 in radish. The distribution of this T6SS cluster in the *S. rhizophila* complex could make it possible to exploit these strains as biocontrol agents against *X. campestris* pv. *campestris*.

## 1 INTRODUCTION

Disease emergence of plant pathogens is the result of changes in host range and/or pathogen dispersion into a new geographic area (Engering et al., 2013). Regarding this second point, seed transmission represents an important means of pathogen dispersion and is therefore significant in the emergence of plant diseases (Baker and Smith, 1966). Indeed, the International Seed Testing Association Reference Pest List v9 identified 333 seed-borne pests (viruses, bacteria, fungi, oomycetes and nematodes) in more than 50 plant species (Denancé and Grimault, 2022). Of these seed-borne pests 145 are directly transmitted to plants (Denancé and Grimault, 2022).

Pathogens are not the only microorganisms that can be carried by seeds. More than 1,000 bacterial and fungal taxa were identified in the seed microbiota of 50 plant species (Simonin et al., 2022). This important microbial diversity observed on seed samples is however more restricted at the scale of an individual seed with one dominant taxon of variable identity (Chesneau et al., 2022; Newcombe et al., 2018). Since the seed is a limited habitat in terms of resources and space, microbial competition is likely to play an important role in seed microbiota assembly. Using these competition processes to promote seed transmission of non-pathogenic microorganisms at the expense of plant pathogens could be deployed as a biocontrol-based strategy (Barret et al., 2016). However, this approach requires a better understanding of the mechanisms involved in these microbial competition processes, notably the relative importance of exploitative competition (i.e. increase uptake and use of nutrients) versus interference competition (i.e. limiting the access of other cells to resources, Granato et al., 2019).

The *Lysobacteraceae* (earlier known as *Xanthomonadaceae*) family includes numerous species of plant pathogens like *Xanthomonas* spp. (Jacques et al., 2016) and also ubiquitous *Stenotrophomonas* spp. like commensal *S*. *rhizophila* (Wolf et al., 2002) and opportunistic human pathogens like *S*. *maltophilia* (Gröschel et al., 2020). *X*. *campestris* pv. *campestris* (Xcc), the causal agent of black rot disease of *Brassicaceae* (Vicente and Holub, 2013) is not only seed-transmitted in a range of *Brassicaceae* (Randhawa, 1984; Rezki et al., 2016; van der Wolf et al., 2019) but also in non-host plants such as common bean (Darrasse et al., 2010). Diversity surveys of the radish seed microbiota have highlighted that Xcc shares the same habitat as numerous bacterial strains related to the *S. rhizophila* species (Rezki et al., 2016, 2018). Strains of *S. rhizophila* are efficient seedling colonizers of cotton, tomato, sweet pepper (Schmidt et al., 2012) and radish (Simonin et al., 2023) as well as commonly isolated in the rhizosphere of different plant species including rapeseed (Berg et al., 1996) and potato (Lottmann et al., 1999). The type strain of *S. rhizophila*, DSM14405^T^, can protect plants against osmotic stress (Alavi et al., 2013; Egamberdieva et al., 2011) and limits the growth of fungal pathogens (Minkwitz and Berg, 2001). Other strains of *S. rhizophila* possess antibacterial activities (Lottmann et al., 1999). For instance, the strain *S. rhizophila* CFBP13503 decreases the population size of Xcc during *in vitro* confrontation assays (Torres-Cortés et al., 2019). This decrease in Xcc population size was attributed to exploitative competition since these strains shared significant overlaps in resource utilization (Torres-Cortés et al., 2019). However, the role of interference competition was only partially assessed through the production of diffusible molecules, while contact-dependent mechanisms were not tested.

Among the contact-dependent mechanisms involved in interbacterial competition, the type VI secretion system (T6SS) is probably the most widely distributed with more than 17,000 T6SS gene clusters distributed in more than 8,000 genomes sequences (source SecReT6 v3, Zhang et al., 2022). T6SS is a multi-protein complex composed of several core components, including the membrane complex TssJLM, the baseplate TssEFGK, the tail tube Hcp (TssD), the spike composed of VgrG (TssI) trimers topped by a protein containing a Pro-Ala-Ala-Arg Repeat (PAAR) motif, the contractile sheath TssBC, and the coordinating protein TssA, as well as the sheath disassembly ATPase ClpV (also known as TssH) (Cherrak et al., 2019; Cianfanelli et al., 2016; Ho et al., 2014). The T6SS allows bacteria to compete and survive in their environments by injecting toxins/effectors into target cells. Effectors are either fused ("specialized" effectors) to or interact ("cargo" effectors) with Hcp tube or VgrG/PAAR spike proteins (Cherrak et al., 2019; Cianfanelli et al., 2016; Jurėnas and Journet, 2021). The contraction of the sheath leads to the injection of Hcp and VgrG/PAAR proteins together with the effectors. Effector-immunity encoding gene pairs are often associated with genes encoding the elements involved in their delivery. These include Hcp, VgrG or PAAR proteins, and also accessory proteins named chaperones/adaptors, which facilitate the loading of effectors onto the T6SS elements (Unterweger et al., 2017). Adaptors identified so far are DUF4123, DUF1795, DUF2169, and DUF2875-containing proteins encoded upstream of their cognate effector (Berni et al., 2019; Unterweger et al., 2017). If effector-immunity can be orphan genes, they are often encoded in T6SS clusters or associated with *hcp* or *vgrG* in orphan *hcp/vgrG*- islands. Conserved domains or motifs have also been described for some effectors and immunity proteins which facilitate their identification (Lien and Lai, 2017). For effector proteins, these conserved domains can reflect their biochemical toxic activity. Recruitment domains and motifs, such as MIX (Marker for type sIX effectors) or FIX (Found in type sIX effectors) motifs, DUF2345/TTR (Transthyretin-like domains) domains or Rhs (Rearrangement hot spot) domains can be found in T6SS effectors and are related to their mode of delivery (Cianfanelli et al., 2016; Cherrak et al., 2019; Jurėnas and Journet, 2021).

Type VI secretion system is widespread among plant-associated bacteria and divided into five taxonomic groups (Bernal et al., 2018). T6SS has been implicated in a wide range of biological processes, including microbial competition with bacteria and fungi (Luo et al., 2023; Trunk et al., 2018), epiphytic colonization of bacterial pathogens (Liyanapathiranage et al., 2021), and pathogen virulence (Choi et al., 2020; Montenegro Benavides et al., 2021; Shyntum et al., 2015). *Pseudomonas putida* KT2440 K1-T6SS also provides biocontrol properties by killing *X. campestris* when injected into plant leaves (Bernal et al., 2017). Commensal bacteria from seed microbiota carrying T6SS could be good candidates as biocontrol agents and may be used to limit bacterial pathogen transmission from seed to seedling. In the course of this work, we explored the possibility of limiting Xcc transmission from seed to seedling through a contact-dependent T6SS mechanism mediated by *S*. *rhizophila*. We also ask to which extent strains from the *Stenotrophomonas* genus share similar or different T6SS clusters.

## 2 RESULTS

### Genomic organization of *S. rhizophila* CFBP13503 T6SS

Fourteen genes encoding T6SS core protein components were identified in the genome sequence of *S. rhizophila* CFBP13503 (**Figure 1**). These genes, located on a single genomic cluster of 72 kb, include genes encoding proteins involved in the membrane complex (TssJ, L and M: HKJBHOBG_02310, 02312, 02279), the baseplate (TssE, F, G and K: HKJBHOBG_02304, 02303, 02302, 02311), the contractile sheath (TssB and C: HKJBHOBG_02307, 02306), the coordinating protein (TssA: HKJBHOBG_02276), the disassembly ATPase (TssH: HKJBHOBG_02301), the inner tube (Hcp: HKJBHOBG_02305) and the puncturing structure (VgrG and PAAR). Regarding the puncturing structure, multiple genes encoding VgrG (n=7) and PAAR domain-containing proteins (n=5) were detected in the genome sequence.

**Figure 1.**
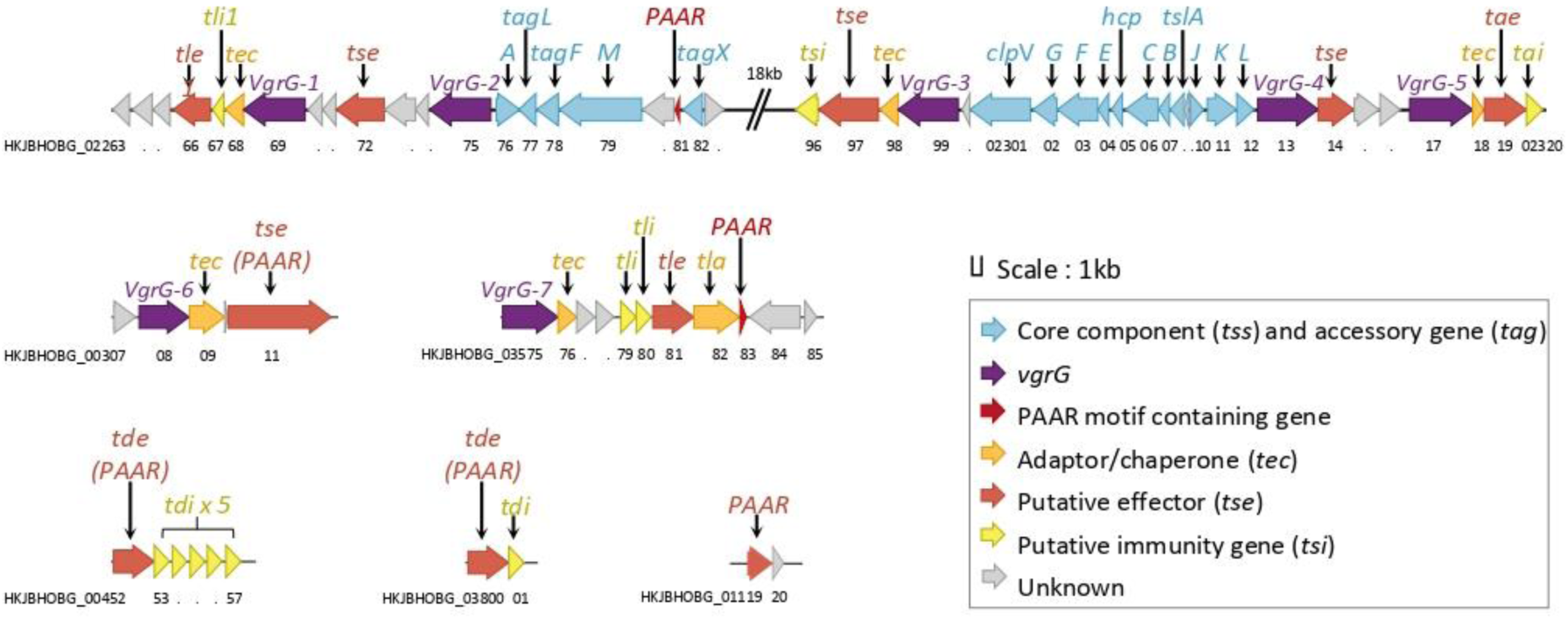
*S. rhizophila* CFBP13503 T6SS genomic architecture. Schematic representation of *S. rhizophila* CFBP13503 T6SS cluster along with orphan *vgrG* and PAAR clusters. Genes are coloured according to their functional roles: core component and accessory genes (blue), *vgrG* (violet), PAAR-motif containing genes (red), adaptor/chaperone (orange), putative effectors (orange-red), immunity genes (yellow), and genes with unknown function (grey).

Based on the genetic architecture of the T6SS genomic cluster, the T6SS is related to group i4 (Bayer-Santos et al., 2019). T6SS putative accessory genes were identified in the core structural cluster, including the group 4-specific : *i)* the group 4-specific *tagX* (HKJBHOBG_02282), which encodes an L,D-endopeptidase first proposed to be involved in cell wall degradation for T6SS assembly (Weber et al., 2016), but essential for polymerisation of the contractile sheath and not required for assembly of the membrane complex and the baseplate (Lin et al., 2022), *ii*) *tagF* (HKJBHOBG_02278), which encodes a negative post-translational regulator of the *P. aeruginosa* H1-T6SS (Lin et al., 2018), *iii*) *tagN/L* (HKJBHOBG_02277), whose role in T6SS assembly or regulation remains unknown and *iiii*) *tslA* (HKJBHOBG_02308), conserved in i4b T6SS and involved in cell-contact T6SS assembly (Lin et al., 2022).

### The T6SS of *S. rhizophila* CFBP13503 encodes a wealth of putative effectors

To identify putative T6SS effectors of CFBP13503, we screened the T6SS main cluster and the *vgrG* and PAAR islands for the presence of "specialized" VgrG, Hcp, or PAAR proteins, as well as the presence of N-terminal effectors motifs such as FIX or MIX domains. We also looked for conserved domains encoded by the genes in the vicinity of *vgrG*, *hcp* or *PAAR* or adaptor/chaperones that could be cargo effectors associated with immunity proteins.

We identified only one Hcp protein, encoded in the main T6SS cluster in between *tssE* and *tssC* genes. Seven *vgrG* genes (*vgrG*-1 to *vgrG*-7) were identified, with five of them located adjacent to the T6SS cluster and two others scattered throughout the genome (**Figures 2a** and **2b**). All VgrG proteins, except VgrG-6, contain a C-terminal DUF2345 domain extending the VgrG needle that can be important for recruiting effectors or carrying toxic activity (Flaugnatti et al., 2016, 2020; Storey et al., 2020). In contrast, VgrG-6 contains a C-terminal domain extension with a weak (*p*-value = 5.52e-03) similarity to the FliC/FljB family flagellin (accession: cl35635). Five PAAR-containing proteins were detected in the genome of CFBP13503: one in the T6SS cluster, one in each vgrG island and two orphans PAAR proteins (**Figure 2**). The latter two PAAR-containing proteins possess a C-terminal Ntox15 domain (pfam15604) with a HxxD catalytic motif. Such domains were found associated with T6SS effectors with DNAse activity called Tde (Type VI DNase effector) (Ma et al, 2014; Luo et al 2023). These two predicted PAAR-fused Tde proteins are associated with genes encoding the DUF1851 domain, known to be associated with T6SS Tde immunity proteins (Tdi) (Ma et al, 2014; Luo et al 2023) (**Figure 2c**).

**Figure 2.**
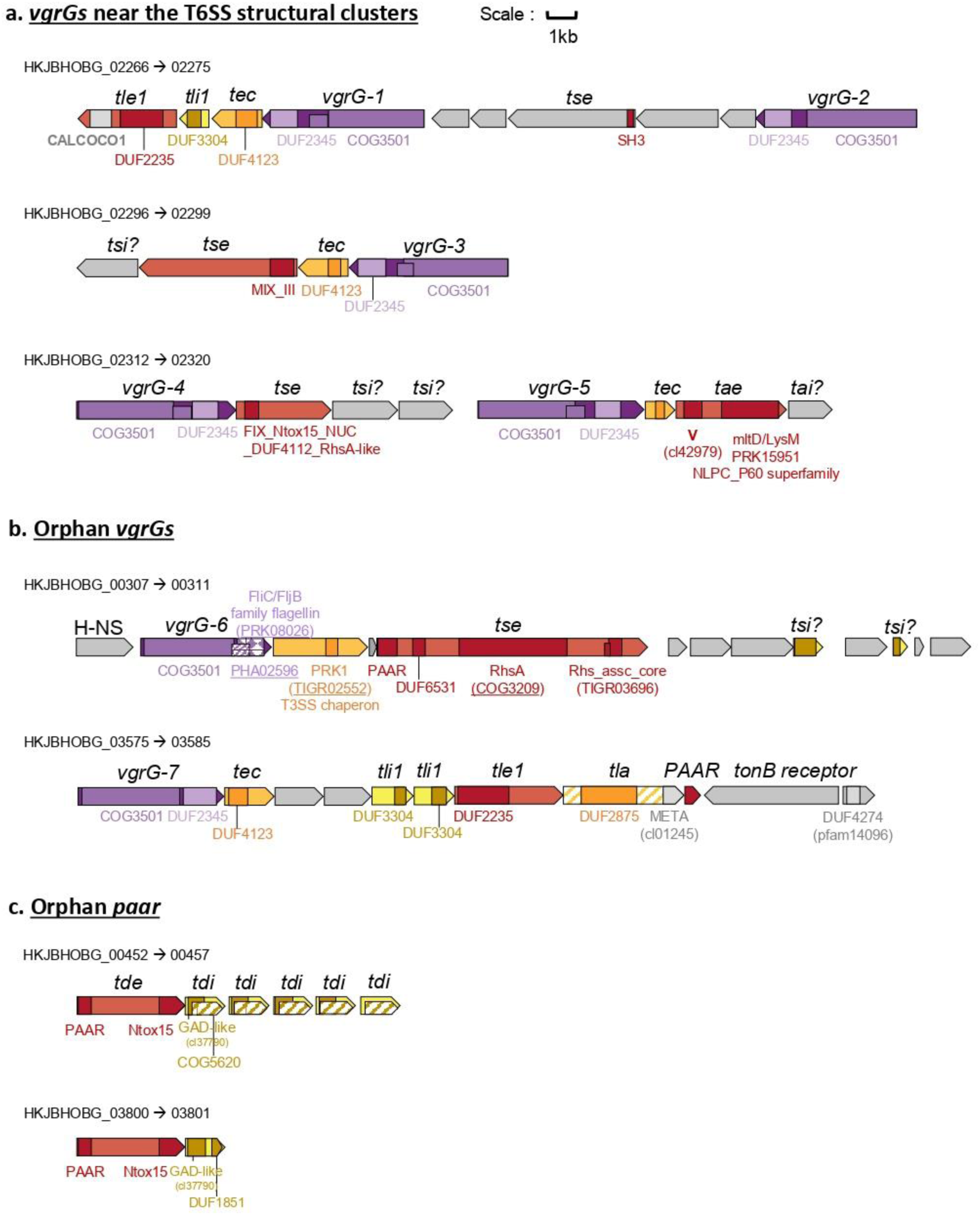
*S. rhizophila* CFBP13503 T6SS effectors. The conserved domain organization of genes encoding putative chaperone, effector and immunity proteins associated with (**a**) VgrG encoded in the main T6SS cluster, (**b**) orphan VgrG, and (**c**) orphan PAAR domain-containing effectors are represented. VgrG genes are indicated in violet, putative effectors with or without PAAR motif are shown in red, chaperone genes in orange, immunity protein genes in yellow and gene-encoded proteins with unknown functions are in grey.

In addition to “specialized” effectors, we detected several putative “cargo” effectors, immunity proteins and chaperone proteins. Regarding chaperones, three DUF4123 domain-containing proteins were identified in CDS located immediately downstream of *vgrG-1, vgrG-3 and vgrG-7*. Another chaperone containing a DUF2875 domain was also encoded in the vicinity of *vgrG-7*. Concerning effector, two effector-immunity (E-I) encoding gene pairs *tle/tli* were associated with VgrG-1 and VgrG-7. Both *tle* encode proteins containing a GXSXG motif (GFSRG) and a DUF2235, characteristic of the T6SS phospholipase effector (Tle) of the Tle1 family. (Flaugnatti et al., 2016; Russell et al., 2013). Their corresponding Tli contained a DUF3304 domain found in several T6SS Tli1 immunity proteins (Russell et al., 2013). A putative amidase effector, *tae*, was associated with VgrG-5. This effector contained a murein transglycosylase D domain and a LysM motif involved in binding peptidoglycan, as well as an NlpC_P60 domain. Linked to the other *vgrG* genes, we also detected genes encoding T6SS effectors specific domains MIX and FIX. The MIX-containing effector located at the vicinity of *vgrG-3* and a tec protein-encoding gene may encode a pore-forming toxin. This toxin is predicted to encode 4-5 putative transmembrane domains at its C-terminus. A predicted structure-based search using Alphafold2 predicted structure and Foldseek suggest that this protein shares structural homologies (over the 900 first residues) with VasX, a pore-forming toxin from *Vibrio cholerae* (Miyata et al., 2013). The downstream gene is a predicted inner membrane protein and could be the corresponding immunity protein. Unfortunately, we were unable to determine conserved domains or potential activities for the FIX-containing protein encoded downstream vgrG-4 so we referred to it as *tse*, which stands for "type six effector".

Similarly, no function could be assigned to the proteins encoded by the genes downstream *vgrg-4*. We assume that these genes could encode potential toxins and associated immunity proteins and potential chaperones. The 18kb inter region between *tssM* and *tssA* and other structural genes contains a poly-immunity locus with a predicted formylglycine-generating enzyme family immunity protein encoding gene (Lopez et al. 2021) and 8 duplications of the putative immunity gene HKJBHOBG_02263 containing a DUF6708 domain.

### *S. rhizophila* CFBP13503 outcompete Xcc8004 *in vitro* in a T6SS-dependent manner

*Stenotrophomonas rhizophila* CFBP13503 is able to outcompete the phytopathogenic bacterial strain Xcc8004 in TSB10 (Torres-Cortés et al., 2019). After 6 hours of confrontation on TSA10 medium between Xcc8004 and CFBP13503, a 10 to 100-fold decrease in Xcc8004 population was observed in comparison to Xcc8004 monoculture (**Figure 3**). Deletion of two genes encoding proteins involved in T6SS assembly (Δ*hcp* and Δ*tssB*) significantly increased the population size of Xcc8004 to a level comparable to Xcc8004 monoculture (**Figure 3**). Complementation of these two mutants restored the decrease in CFU of Xcc8004. These results demonstrate that the T6SS of *S. rhizophila* CFBP13503 is involved in the antibiosis towards Xcc8004 from 6 hours of confrontation. T6SS effect size increased with time as 24 hours after confrontation Xcc8004 population decreased from 1,000 to 10,000 times in the wild-type strain and complemented mutants. At 48h the Xcc populations were reduced by the wild type compared to the deletion mutants but increased by one log10 compared to the same wild type treatment at 6 and 24 h (**Fig. S3**). These results highlight the involvement of the T6SS of *S. rhizophila* CFBP13503 in the killing of Xcc8004 and thus its involvement in interbacterial competitions.

**Figure 3.**
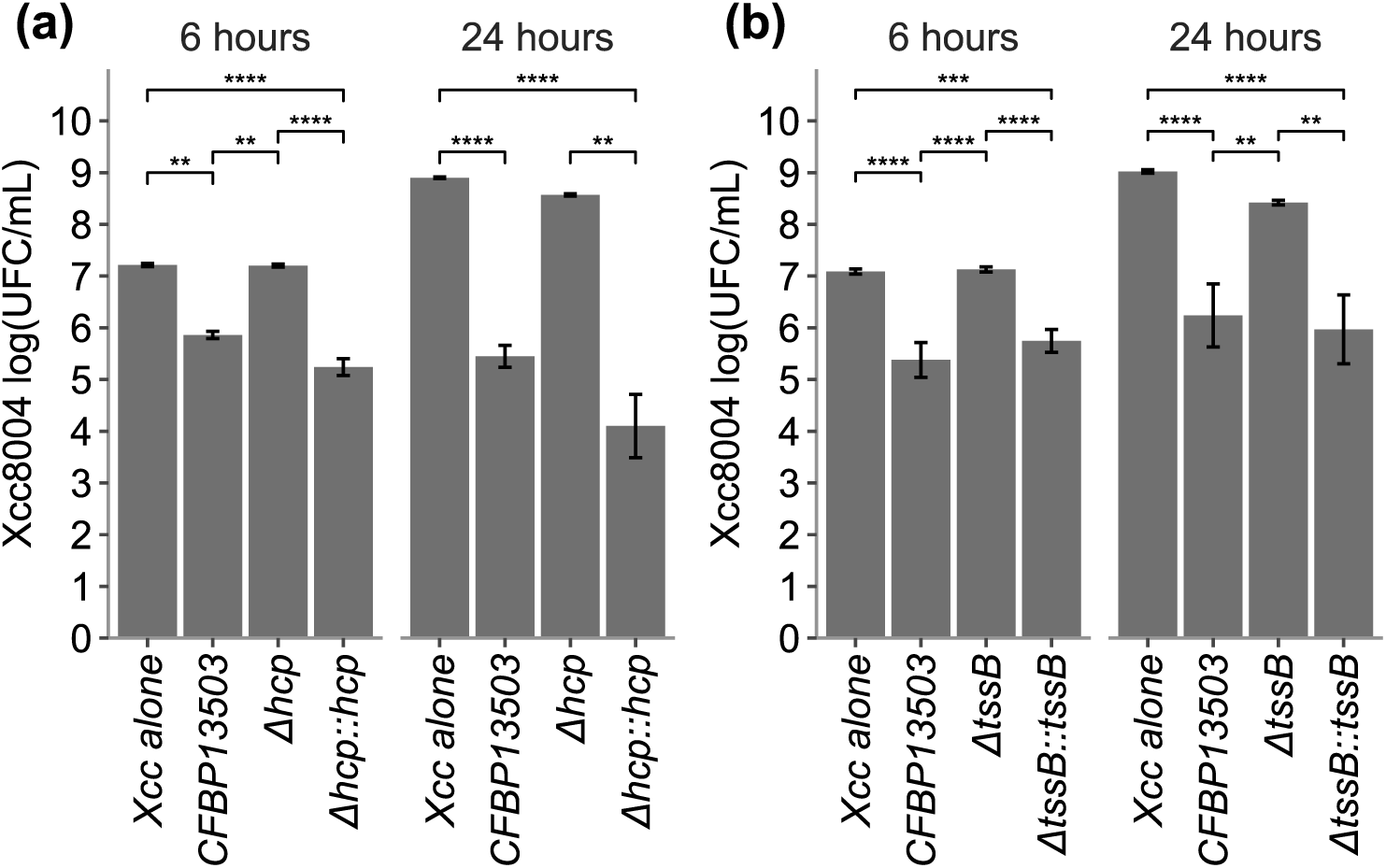
Bactericidal activity of *S. rhizophila* CFBP13503 against *X. campestris* pv. *campestris* 8004. Confrontation between Xcc8004 and i) *S. rhizophila* CFBP13503 wild-type, ii) T6SS-deficient mutants *Δhcp* (**a**) and *ΔtssB* (**b**) and iii) the complemented mutants *Δhcp::hcp* and *ΔtssB::tssB* in TSA10 medium for 6h and 24h. Colony-forming units (CFU) were quantified on TSA10 supplemented with rifampicin. The averages ± sd of 9 replicates are plotted. Statistical analyses were performed using Dunn’s Multiple Comparison Test (* p-value < 0,05; ** p-value < 0,005; *** p-value < 0,0005; **** p-value < 0,00005).

### The T6SS of *S. rhizophila* CFBP13503 limits the seed-to-seedling transmission of Xcc8004 in radish

To investigate the bactericidal impact of CFBP13503 T6SS *in planta*, seedling transmission assays were carried out on radish seeds (**Figure 4**). Xcc8004 was co-inoculated with either the wild-type CFBP13503 strain or the *Δhcp* mutant on sterilized seeds with inoculum ratios of 1:2.1 and 1:1.6 respectively. Hcp protein is a structural protein of T6SS syringe but could exerts antibiosis against bacterial preys (Decoin et al. 2014, Fei et al. 2022). Thus, we chose the *hcp* mutant to prevent any toxicity. Bacterial population sizes were measured on seeds, germinated seeds (1 dpi) and the aerial and root parts of seedlings (5 dpi) by quantification of CFU on selective media. The Xcc8004 population of inoculated seeds presented a variable (6.1 to 6.5 log_10_(UFC/sample)) but no significant decrease when co-inoculated with wild-type CFBP13503 compared to the *Δhcp* mutant. However, a significant decrease in the Xcc8004 population was observed from the germinated seed stage when co-inoculated with the wild-type strain. This decrease persisted over time, with a significant reduction in Xcc8004 population size in the aerial and root part of seedlings during the confrontation with the wild-type CFBP13503 (**Figure 4**). The population size of *S. rhizophila* was not impacted by the presence or absence of a functional T6SS (**Figure 4**). Altogether these findings show that the T6SS of *S. rhizophila* CFBP13503 restricted seedling transmission of Xcc8004 without providing a fitness advantage to *S. rhizophila* CFBP13503.

**Figure 4.**
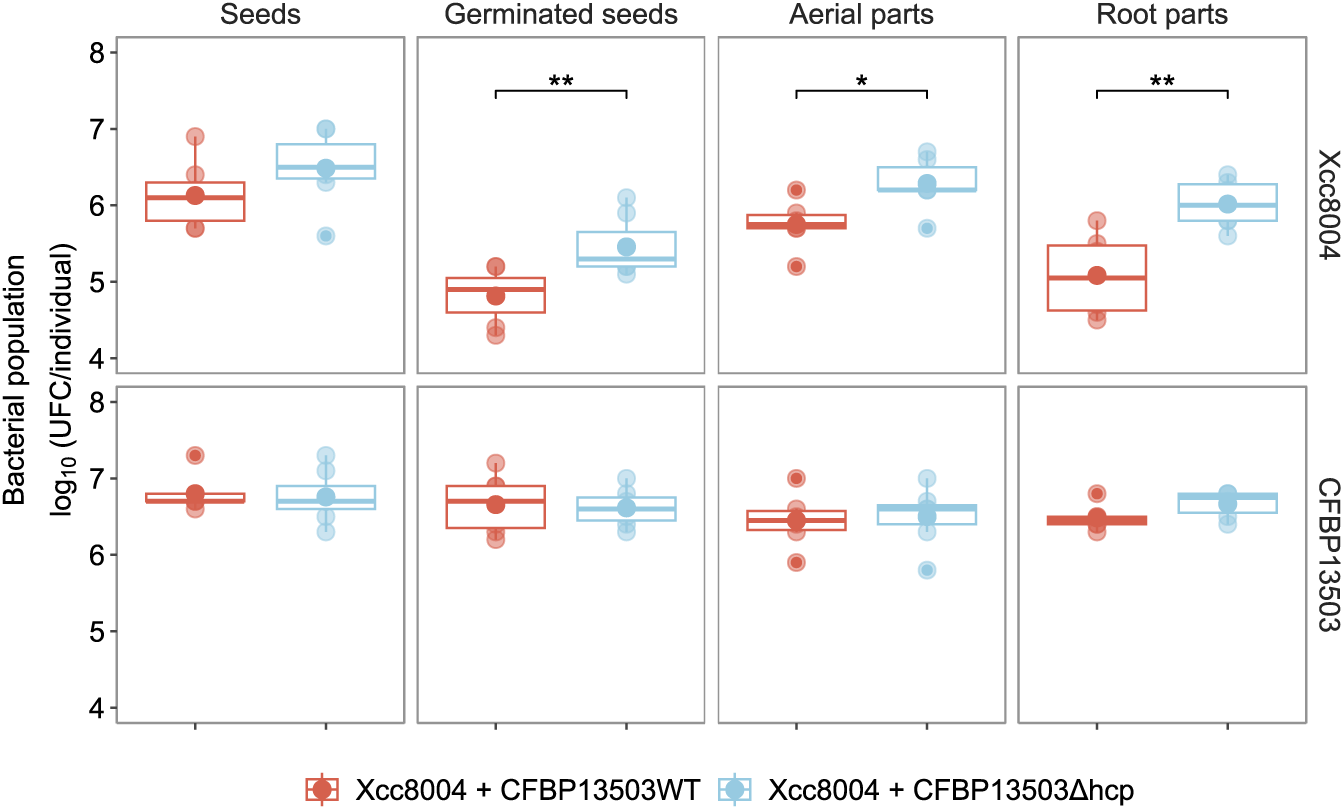
Bacterial population dynamics during seed-to-seedling transmission of *S. rhizophila* CFBP13503 and Xcc8004 in radish. Radish seeds were co-inoculated with Xcc8004 and (i) CFBP13503 wild-type or (ii) T6SS-deficient *Δhcp*. Bacterial populations were assessed in radish seeds (0 dpi), 24-hour-old germinated seeds (1 dpi), 5-day-old seedling aerial parts, and 5-day-old seedling root parts (5 dpi). Colony-forming units (CFU) were quantified on selective media to differentiate Xcc8004 and CFBP13503 populations. The averages ± sd of two experiments (n=3 and n=4) are plotted. Statistical analyses were performed using Dunn’s Multiple Comparison Test (* p-value < 0,05; ** p-value < 0,005).

### Analysis of T6SS distribution within the *Stenotrophomonas* genus

Given the significant effect of *S. rhizophila* CFBP13503 T6SS on Xcc8004, it raises the question of T6SS distribution within the *Stenotrophomonas* genus. A total of 835 *Stenotrophomonas* genome sequences were then collected from the NCBI (experimental procedure). These genome sequences are divided into 95 groups (**Figure S1**) at a threshold of 50% shared 15-mers, an overall genome relatedness index employed as a proxy for species delineation (Briand et al., 2021). From this analysis, the strain CFBP13503 is grouped with *S. rhizophila* DSM14405^T^ in the same species complex (threshold 0.25) but differs from the DSM14405^T^ type strain of *S*. *rhizophila* (threshold 0.5). So, we describe here a new *S*. *rhizophila* complex.

Sixty-four strains (8.3% of the dataset) of 22 groups contained at least one T6SS cluster (**Figure 5**). Based on the phylogenetic analysis of TssC (**Figure 6**), T6SS clusters were classified into three families i1 (*n*=5), i3 (*n*=10), and i4 (*n*=57). Just four species among the 28 groups of *S*. *maltophilia* species complex contain one T6SS except group 65 which contains two T6SS belonging to two different taxonomic groups (i3 and i4). All *S. rhizophila* strains (**n=5**) have a T6SS related to group i4.

**Figure 5.**
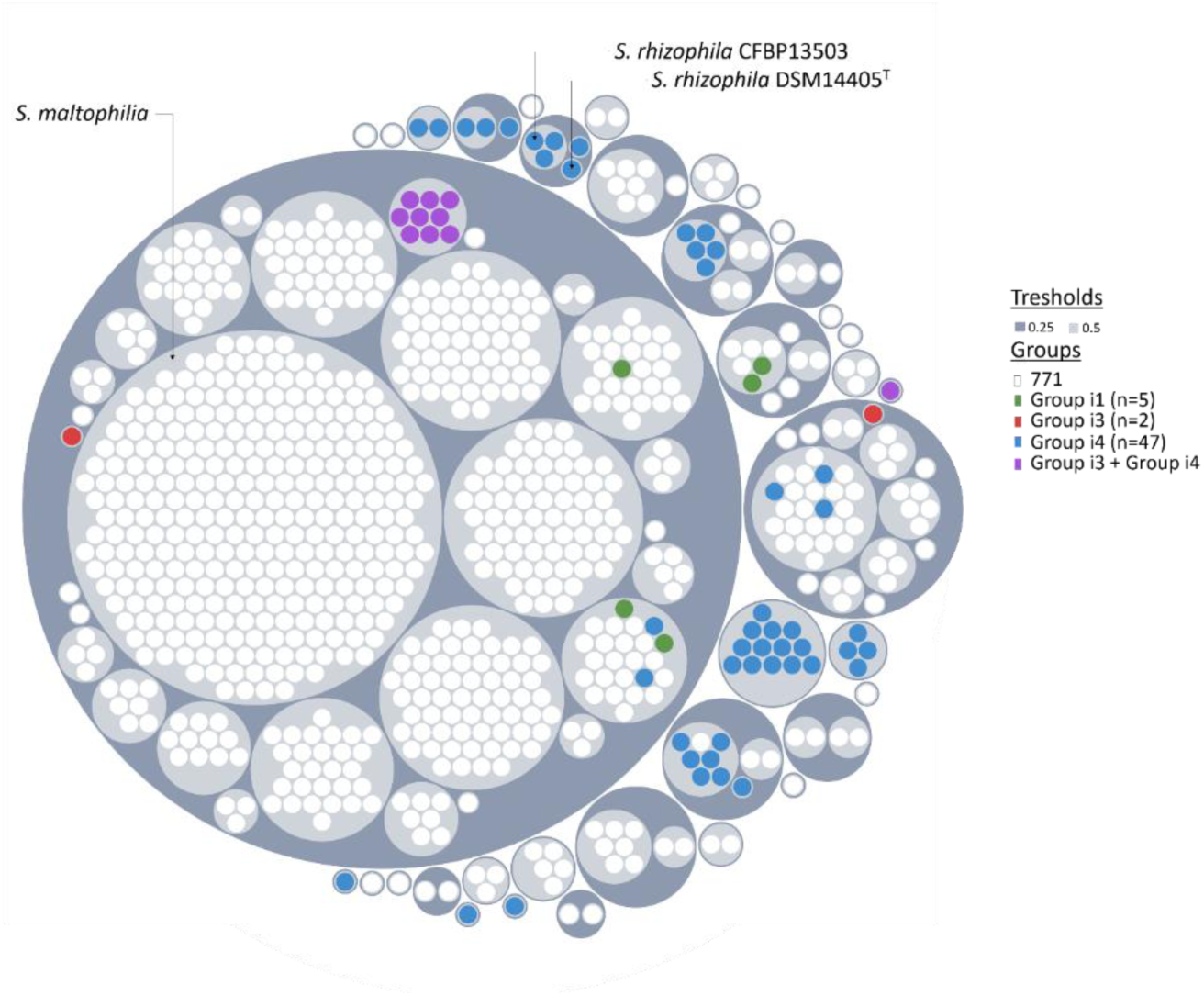
T6SS distribution among *Stenotrophomonas* sp.. Circle packing representation of *Stenotrophomonas* sp. genomes (n=835). Overall genome relatedness was assessed by comparing the percentage of shared 15-mers. Each dot represents a genome sequence, color-coded based on the T6SS group. The genomes were grouped using two distinct thresholds to assess species-specific relationships (0.5) and interspecies relationships (0.25). Interactive circle packing representation is available in Figure S1. For the caption of species names when a reference strain exists see Figure S2.

**Figure 6.**
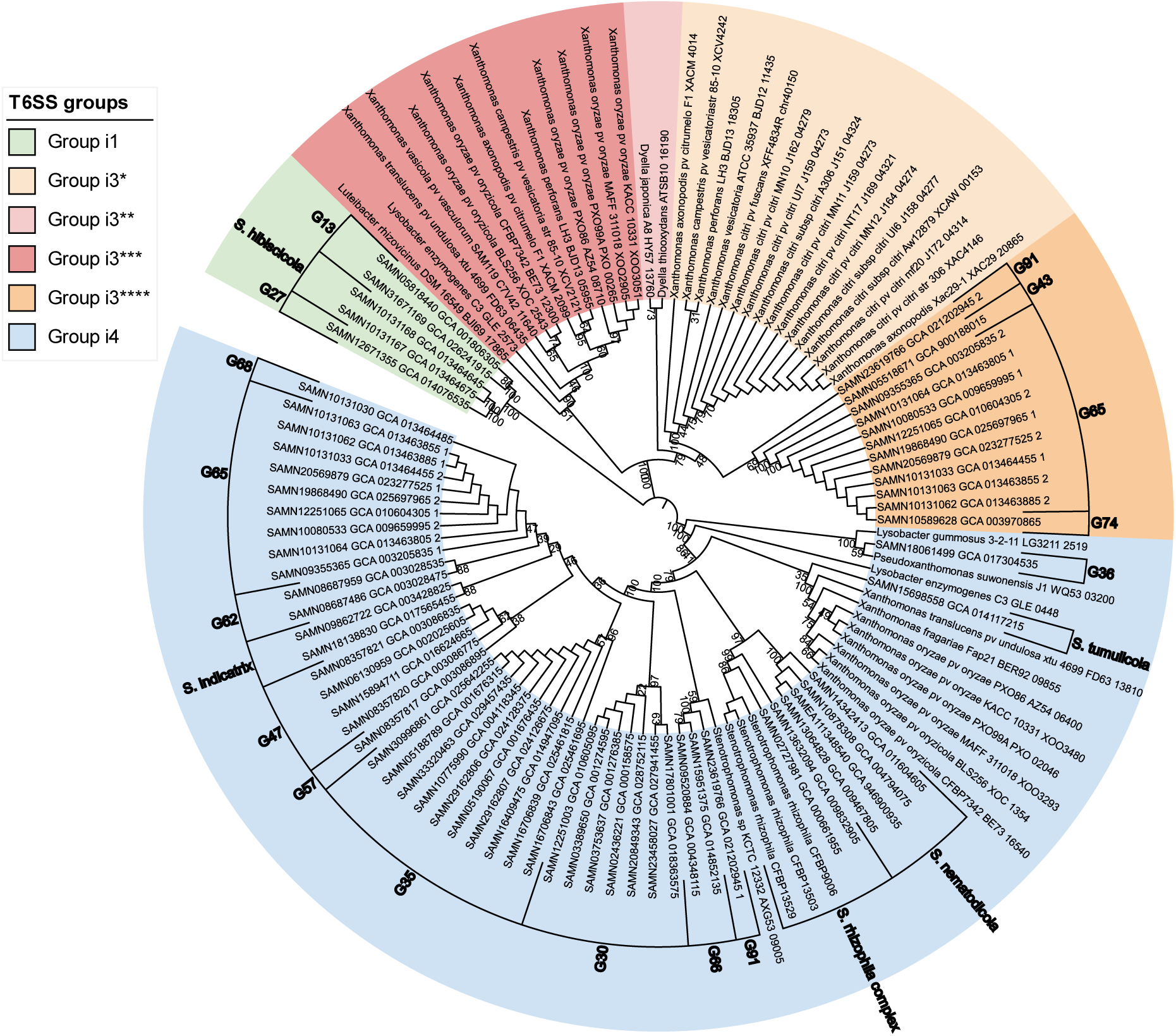
T6SS distribution among *Lysobacteraceae*. Phylogenetic tree based on TssC protein sequences, constructed using a Kimura two-parameter neighbor-joining method with 1000 bootstrap replicates. The tree includes *Stenotrophomonas* and other *Lysobacteraceae*, as studied by Bayer-Santos et al. in 2019. Strain taxonomic affiliation is based on KI-S grouping (Figure S1).

## 3 DISCUSSION

*Xanthomonas campestris* pv. *campestris* is a frequent seed-colonizer of various host and non-host plant species (Darrasse et al., 2010; Vicente and Holub, 2013). In the seed habitat, Xcc can co-occur with other bacterial species such as *S. rhizophila* CFBP13503 (Rezki et al., 2016), which can outcompete Xcc8004 *in vitro* (Torres-Cortés et al., 2019). Resource overlap was initially proposed as the mode of action involved in Xcc8004 growth reduction in the presence of *S. rhizophila* CFBP13503 (Torres-Cortés et al., 2019).

In this study, we showed that the growth inhibition of Xcc8004 by *S. rhizophila* CFBP13503 was T6SS-dependent. CFBP13503 decreased Xcc8004 population size from 10 to 1,000-fold after 6h and 24h of confrontation in solid medium, respectively. Deletion mutants of two genes encoding proteins (Δ*hcp* and Δ*tssB*) involved in T6SS assembly were no longer able to reduce Xcc8004 growth. After 48h of confrontation *in vitro*, Xcc8004 strain partially escaped T6SS antibiosis due to the possible establishment of non-immune defenses by Xcc such as the creation of dead cell barriers or the production of exopolysaccharides that limit cell-to-cell contact (Hersch et al., 2020). Antibacterial activities of T6SS can be contact-dependent with the translocation of effector proteins in bacterial cells (Jurėnas and Journet, 2021) or contact-independent with the secretion of metal scavenging proteins in the surrounding media (Chen et al., 2016; Lin et al., 2017; Si et al., 2017). When the confrontation between Xcc8004 and CFBP13503 took place in a liquid medium with limited cell contacts, no difference in Xcc8004 growth was observed between the wild-type strain and the T6SS-deficient mutants (**Fig. S4**). In conclusion, T6SS-mediated reduction in Xcc8004 growth is contact-dependent and therefore due to the injection of protein effectors in Xcc8004 cells.

Furthermore, *S. rhizophila* CFBP13503 is involved in reducing the transmission of Xcc8004 from seed to seedling. When comparing equivalent Xcc8004 populations on seeds, the presence of *S. rhizophila* CFBP13503 negatively impacted the population of Xcc8004 on germinated seeds compared to the mutant strain lacking the T6SS gene *hcp*. This highlights the potential role of T6SS during the early interactions between bacteria. The effects of *S. rhizophila* CFBP13503 T6SS persist overtime at the seedling stage, as a lower population of Xcc is observed in the presence of the wild-type strain. Since Xcc8004 is a seed-borne pathogen, limiting its transmission to the seedling stage appears to be a promising strategy for managing this pathogen. However, further studies are needed on host plants to assess whether the T6SS of *S. rhizophila* CFBP13503 can limit the pathogenicity of Xcc8004 by reducing its population size.

It is also noteworthy that the presence or absence of T6SS, as well as its mutation, does not impact the transmission of *S. rhizophila* CFBP13503 when in competition with Xcc8004. Consequently, the T6SS of *S. rhizophila* CFBP13503 does not seem to be involved in its adhesion or colonization capacities of radish seed and seedling, unlike what has been observed in other bacterial species (Cassan et al., 2021; Mosquito et al., 2019).

*S. rhizophila* CFBP13503 possesses seven VgrG proteins, each of which is associated with a chaperone protein and a putative effector. This diversity of VgrG proteins may allow for various associations in the arrangement of VgrG trimers, enhancing the versatility of the T6SS.

Additionally, *S. rhizophila* CFBP13503 has five PAAR-domains containing protein. The presence of PAAR proteins sharpens the tip of the VgrG trimer, creating opportunities for different toxic effector associations during each firing event. Two of these PAAR proteins possess a C-terminal toxic domain related to DNase activity. The extensive repertoire of effectors identified in *S. rhizophila* CFBP13503 includes Tle1-type phospholipases (Flaugnatti et al., 2016; Russell et al., 2013) capable of lysing the membranes of target bacteria, DNases (Tde) that exhibit antibacterial properties by targeting nucleic acids, amidases (Tae) that degrade peptidoglycan to lyse the target bacterium, and potentially a pore-forming effector that inhibits the growth of target cells by depolarizing the inner membrane. Furthermore, there are other effectors whose activities have yet to be discovered. These effectors are predicted to target different components of bacterial cells, contributing to the antibacterial phenotype against Xcc8004. Interestingly, some components and effectors in *S. rhizophila* CFBP13503 could also exhibit anti-eukaryotic activity. For example, the DUF2345 domain of VgrG has been shown to intoxicate yeast, as demonstrated by *Klebsiella pneumoniae* VgrG4 (Storey et al., 2020). Moreover, the two "evolved" PAAR proteins, which contain a C-terminal Ntox15 domain, share a significant identity (>30%) with the DNAse effector TafE of *Acinetobacter baumannii* strain 17978, known for its involvement in yeast killing (Luo et al., 2023). Some Tle effectors were also shown to target bacteria and eukaryotic cells (Jiang et al., 2014; Jiang et al., 2016). The combination of these diverse effectors, either individually or in synergy (LaCourse et al., 2018), likely contributes to the competitive advantage of *S. rhizophila* CFBP13503 over Xcc *in vitro*. Despite possessing only one T6SS, the diverse repertoire of effectors enables *S. rhizophila* CFBP13503 to effectively combat bacterial competitors and potentially exert antibacterial effects against Xcc8004.

Type VI secretion system is not frequently found in *Stenotrophomonas* genomes. Less than 10% of the genome sequences analyzed possessed a T6SS genetic cluster, which is generally presented in a single copy. This frequency is relatively low compared with other bacterial genera of the *Lysobacterales* where T6SS is present in approximately 50% of sequenced strains (Bayer-Santos et al., 2019). A novel subgroup 3 within *Lysobacterales*, exclusively associated with *Stenotrophomonas* species, was revealed by TssC-based phylogeny analysis. Notably, this subgroup was not identified in the previous study conducted by Bayer-Santos et al. (2019) on T6SS classification within *Lysobacterales*. Consistent with their findings, our analysis also identified *Stenotrophomonas* T6SS classified into both group i1 and group i4. However, group i4 T6SS appears to be more prevalent within the *Stenotrophomonas* genus. This is the case for the T6SS of *S. rhizophila* CFBP13503 and more generally for strains affiliated with the *S. rhizophila* complex, including the type strain DSM14405^T^ (Wolf et al., 2002). *S. rhizophila* strains are not only known for their ability to colonize a wide range of plant species following seed inoculation (Schmidt et al., 2012; Simonin et al., 2023) but also to exhibit antifungal (Berg and Ballin, 1994) and antibacterial activities (Lottmann et al., 1999). For instance, the strain *S. rhizophila* DSM14405^T^ protects plants against *Fusarium solani* and displays antagonistic activity against various phytopathogenic fungi under high salt conditions (Egamberdieva et al., 2011). Interestingly some T6SS genes of *S. rhizophila* DSM14405^T^ are strongly up-regulated in response to osmotic stress (Alavi et al., 2013; Liu et al., 2022). More available genomes from the *S*. *rhizophila* complex will confirm later the T6SS uniformity in these species. Nevertheless, the large range of putative T6SS effectors in CFBP 13503 reinforces the interest in *S*. *rhizophila* antimicrobial activities.

The T6SS of *S. rhizophila* CFBP13503 plays a crucial role in its antibiosis against Xcc8004 and in limiting Xcc8004 transmission from radish seed to seedling, highlighting its potential in biocontrol of seed-borne pathogenic bacteria. The T6SS has emerged as a powerful tool in biocontrol strategies, offering a novel approach to combat plant pathogens. However, further research is necessary to fully understand its impact within complex microbial ecosystems. By investigating the role of the T6SS in diverse bacterial communities, valuable insights can be gained regarding its functionality, interactions with other microorganisms, and ecological consequences. Understanding the influence of the T6SS on complex bacterial communities is essential for unlocking its full potential and maximizing its contribution to biocontrol approaches.

## 4 EXPERIMENTAL PROCEDURES

### Bacterial strains and growth conditions

Bacterial strains and plasmids used in this study are listed in Table S1. *Stenotrophomonas rhizophila* CFBP13503 and *Xcc* 8004::GUS-GFP (Cerutti et al., 2017) were grown at 28°C on tryptic soy agar 1/10 strength (TSA10: 17 g.l-1 tryptone, 3 g.l-1 soybean peptone, 2.5 g.l-1 glucose, 5 g.l-1 NaCl, 5 g.l-1 K2HPO4, and 15 g.l-1 agar) or tryptic soy broth 1/10 strength (TSB10). *E. coli* DH5α and *E. coli* MFDpir (Ferrières et al., 2010) were grown at 37°C on Luria-Bertani (LB 10 g.l-1 tryptone, 5 g.l-1 Yeast extract, 10 g.l-1 NaCl) medium. LB medium was supplemented with 0.3mM 2,5-diaminopimelic acid (Fisher Scientific, UK) for auxotrophic *E. coli* MFDpir.

### Construction of *S. rhizophila* CFBP13503Δ*hcp* and CFBP13503Δ*tssB* and their complementation

Unmarked *hcp* (HKJBHOBG_02305) and *tssB* (HKJBHOBG_02307) deletions were performed by allelic exchange using the suicide vector pEX18Tc (Hoang et al. 1998). The deletion plasmids pEX18Tc-*Δhcp* and pEX18Tc-Δ*tssB* were constructed using the TEDA cloning procedure (Xia et al., 2019). Briefly, pEX18Tc was digested with *Xba*I (New England Biolabs, France) followed by a dephosphorylation step using the shrimp alkaline phosphatase (Phusion High-fidelity DNA polymerase, New England Biolabs, France). *hcp* and *tssB* flanking regions were PCR-amplified from CFBP13503 with the Phusion High-Fidelity DNA polymerase (New England Biolabs, France), and the primer pairs listed in Table S2. The dephosphorylated pEX18Tc vector and PCR products were purified using the NucleoSpin Gel and PCR Clean-up kit (Macherey-Nagel, Düren, Germany). TEDA reaction was then carried out by mixing 150 ng of pEX18Tc with the corresponding PCR products at a molar ratio of 1:4. One hundred µL of *E. coli* DH5α were transformed with 5 µL of TEDA reaction using the Inoue transformation procedure (Sambrook and Russell, 2006) modified by Xia et al. (2019). Amplicon insertions were validated by colony PCR with the primer pair M13F/M13R. Plasmids (pEX18Tc-*Δhcp*) were extracted with the NucleoSpin plasmid kit (Macherey-Nagel, Düren, Germany), and insertion regions were verified by sequencing (Azenta Life Sciences, Germany). *E. coli* MFD*pir* was transformed with pEX18Tc-*Δhcp* and pEX18Tc-*ΔtssB* using the modified Inoue method. Plasmids were transferred to *S. rhizophila* CFBP13503 *via* conjugation. *S. rhizophila* CFBP13503 transconjugants were selected on TSA10 supplemented with tetracycline (20 µg/mL). The resulting colonies were grown in TSB10 (28°C, 120 rpm, 3h) and bacterial suspensions were spread on TSA10 supplemented with 5% saccharose. Allelic exchanges were validated by PCR and sequencing.

Deletion mutants were complemented by allele exchange. Briefly, *hcp* and *tssB* were PCR-amplified with the primer pairs Hcp-pEX18-UpF/Hcp-pEX18-DnR and TssB-pEX18- UcpF/TssB-pEX18-DnR (Table S2). TEDA reactions were performed at a vector: insert molar ratio of 1:1. pEX18Tc-*hcp* and pEX18Tc-*tssB* were transferred by electroporation to CFBP13503Δ*hcp* and CFBP13503Δ*tssB*, respectively. To carry out this electroporation step, bacteria were grown in TSA to OD_600_ of 0.5-0.7. Bacterial cultures were centrifuged (4100 g, 10 min, 2°C), and pellets were washed four times in cold sterile water, once in 10% glycerol and finally resuspended in 10% glucose before storage at -80°C. CFBP13503Δ*hcp* and CFBP13503Δ*tssB* were transformed with 150 ng of pEX18Tc-*hcp* and *pEX*18Tc-*tssB* (2kV, 5ms). Transformants were selected on TSA10 supplemented with tetracycline (20 µg/mL). The resulting colonies were grown in TSB10 (28°C, 120 rpm, 3h) and bacterial suspensions were spread on TSA10 supplemented with 5% saccharose. Allelic exchanges were validated by PCR and sequencing.

### *In vitro* confrontation assays

*S. rhizophila* CFBP13503, the isogenic T6SS-deficient mutants, the complemented T6SS mutants and Xcc8004-Rif^R^ were cultured overnight in 10 mL of TSB10 (28°C, 150 rpm). Cultures were centrifuged (4,000 g, 8 min, 20°C) and the resulting pellets were resuspended in sterile water. Bacterial suspensions were calibrated to OD_600_ of 0.5 (∼10^9^ cells.mL^-1^). For confrontation on a solid medium, calibrated suspensions were mixed at a ratio of 1:1 (i.e. 100µL of each strain). Single suspensions were prepared as a control by mixing 100µL of bacterial cultures with 100µL of sterile water. Drops of 20 µl were deposited on TSA10, dried for 15 minutes under a laminar and incubated at 28°C for 6 and 24 h. At each incubation time, cells were resuspended in 2.5 mL of sterile water, serial-diluted and plated on TSA10 supplemented with 50 µg/mL of rifampicin (selection of Xcc8004-Rif^R^) or with 50 µg/mL of spectinomycin and 100 µg/mL of ampicillin (selection of *S. rhizophila* strains). For confrontation in liquid medium, calibrated suspensions were mixed at a ratio of 1:1 (i.e. 500 µL of each strain in 9 mL TSB10). As a control, single-strain suspensions were prepared by mixing 500 µL of bacterial cultures with 500 µl of TSB10. The confrontations were incubated at 28°C 150 rpm for 6 and 24h. At each incubation time, confrontations were serial-diluted and plated on TSA10 supplemented with 50 µg/mL of rifampicin (selection of Xcc8004-Rif^R^) or with 50 µg/mL of spectinomycin and 100 µg/mL of ampicillin (selection of *S. rhizophila* strains).

### *In planta* transmission assays

Three subsamples of 300 radish seeds (*Raphanus sativus* var. Flamboyant 5) were surface sterilized using the protocol described in Simonin et al. (2023). Sterilized seeds were dried 30 min before inoculation under a laminar. Bacterial suspensions were prepared at an OD_600_ of 0.5 from 24h bacterial mats on TSA10. These suspensions corresponded to (i) Xcc8004, (ii) Xcc8004/CFBP13503 (1:1 ratio) and (iii) Xcc8004/CFBP13503Δ*hcp* (1:1 ratio). Seeds were either soaked into bacterial suspensions (15 min, 20°C, 70 rpm) or sterile water (non-inoculated condition). Seeds were dried for 15 min on sterile paper under a laminar. Inoculated and non-inoculated seeds were placed on sterile folded moistened papers in sterile plastic boxes. Three repetitions (20 seeds per repetition) were carried out per condition. Boxes were incubated in a growth chamber (photoperiod: 16h/8h, temperature 20°C). Germinated seeds were collected 24 h post-inoculation (n=3, 20 germinated seeds per repetition). Seedlings were harvested five days post-inoculation (n=3, 20 seedlings per repetition). The same experiment was repeated with n=4 repetitions. Bacterial population sizes were assessed by dilution and plating on TSA10 supplemented with appropriate antibiotics (see *in vitro* confrontation assays). Seed-associated bacteria were recovered 15 min after inoculation (initial time) by vortexing 20 seed pools in 2 mL of sterile water for 30 seconds. Germinated seeds (20 seedling pools) were grounded in 4 mL of sterile water. The entire aerial and root parts of seedlings (20 seedling pools) including both endophytic and epiphytic bacteria were separated and grounded in 4 mL of sterile water. No bacterial growth was observed on the selective media for the non-inoculated seeds, germinated seeds and seedlings attesting the absence of culturable bacteria in the control.

### Genomic analysis of *S. rhizophila* CFBP13503 T6SS and effector prediction

The genomic sequence of *S. rhizophila* CFBP13503 (SAMN09062466) was initially obtained through paired-end Illumina sequencing (Torres-Cortés et al., 2019). To circularize the genomic sequence of CFBP13503, PacBio sequencing was performed on an RS2 machine (Genotoul, Castanet-Tolosan, France). PacBio reads were filtered and demultiplexed using the ccs v6.3.0 and lima v2.5.1 tools of the PacBio SMRT Tools v11.0.0.146107 toolkit and then assembled and circularized using Flye v2.9 (Kolmogorov et al., 2019). The sequence start was fixed using the fixstart option of Circlator v1.5.1 (Hunt et al., 2015). Polishing with PacBio reads was performed using Flye v2.9 (Kolmogorov et al., 2019). Polishing with Illumina HiSeq3000 short reads was done using Pilon v1.24 (Walker et al., 2014) with the setting --mindepth 0. Genome annotation was performed with Prokka v1.14.6 (Seemann, 2014).

T6SS components were identified by conducting NCBI BLASTP analysis on protein sequences. Effector-immunity encoding gene pairs and chaperones were identified by analyzing genes downstream of *vgrG* and PAAR motif-containing genes. The conserved domain database of the NCBI was used to identify T6SS-related conserved domains. Structural homology-based searches were made using Alphafold2 (Jumper et al., 2021) structure prediction of putative effectors followed by a DALI (Holm et al., 2023) search analysis or using the Foldseek search server (van Kempen et al., 2023).

### Phylogenetic analysis of T6SS

A total of 991 genome sequences of *Stenotrophomonas* were downloaded from the NCBI. Genomes with fewer than 10 markers (Marker_lineage = f Xanthomonadaceae) absent or in multicopy (CheckM v1.1.6; Parks et al., 2015) were conserved for further analysis. A multi-locus species tree was created with Automlst (Alanjary et al., 2019). Non-*Stenotrophomonas*- affiliated genomes were excluded from further analysis, resulting in a final dataset comprising 835 *Stenotrophomonas* genomes. Sequence relatedness between the selected 835 genomes sequences was assessed with KI-S (Briand et al., 2021) using 50% of shared 15-mers, a proxy for delineating bacterial species. Genome sequences were annotated with Prokka 1.14.6 (Seemann, 2014). The presence of T6SS genomic clusters was predicted with MacSyFinder2 (Néron et al., 2023). TssC protein sequences were retrieved following BLASTp searches using TssC sequences of *Stenotrophomonas rhizophila* CFBP13503 (iT6SS group 4b) and *Stenotrophomonas sp.* LM091 (iT6SS group 1). BLASTp hits with >25% identity over 75% of protein length were conserved. TssC sequences were aligned using MUSCLE. A Kimura two-parameter neighbor-joining tree was constructed with 1000 bootstraps with SeaView v4.7 (Gouy et al., 2010). The T6SS groups have been assigned according to the previously defined nomenclatures (Bayer-Santos et al., 2019; Bernal et al., 2017; Gallegos-Monterrosa and Coulthurst, 2021).

## ACKNOWLEDGEMENTS

This work was funded by the TypeSEEDS project (INRAE: Plant Health and Environment division, HoloFlux Metaprogram) and the Region des Pays de la Loire (RFI “Objectif Végétal”). We thank the CIRM-CFBP for providing access to strains of their culture collection.

## DATA AVAILABILITY STATEMENT

*Sequence data generated during this work can be found in the GenBank database. The coding sequence of CFBP13503 has been deposited in GenBank under accession number CP128598*.

## SUPPORTING INFORMATION LEGENDS

**Figure S1. T6SS distribution among *Stenotrophomonas* sp..** Interactive circle packing representation of *Stenotrophomonas* sp. genomes (n=835). Overall genome relatedness was assessed by comparing the percentage of shared 15-mers. Each dot represents a genome sequence, color-coded based on the T6SS group. The genomes were grouped using two distinct thresholds to assess species-specific relationships (0.5) and interspecies relationships (0.25).

**Figure S2. Clustering of *Stenotrophomonas* genome sequences.** Circle packing representation of *Stenotrophomonas* sp. genomes (n=835). Overall genome relatedness was assessed by comparing the percentage of shared 15-mers. Each dot represents a genome sequence. The coloured dots represent the type strains of each described *Stenotrophomonas* species. The genomes were grouped using two distinct thresholds to assess species-specific relationships (0.5) and interspecies relationships (0.25).

**Figure S3. Dynamics of Xcc8004 and CFBP13503 populations in confrontation from 0h to 48h.** Confrontation between Xcc8004 and i) *S. rhizophila* CFBP13503 wild-type, ii) T6SS-deficient mutants *Δhcp* (**a**) and *ΔtssB* (**b**) in TSA10 medium for 6h, 24h and 48h. Colony-forming units (CFU) of Xcc and CFBP13503 populations were quantified on TSA10 supplemented with rifampicin and ampicillin-streptomycin respectively. The averages ± sd of 6 replicates are plotted. Statistical analyses were performed using Dunn’s Multiple Comparison Test (* p-value < 0,05; ** p-value <0,005; *** p-value < 0,0005; **** p-value < 0,00005).

**Figure S4. Bactericidal activity of *S. rhizophila* CFBP13503 against *X. campestris* pv. *campestris* 8004 in liquid medium.** Confrontation between Xcc8004 and *S. rhizophila* CFBP13503 wild-type, or T6SS-deficient mutants (Δ*hcp* and Δ*tssB*) in TSB10 (liquid medium) for 6h and 24h. Colony-forming units (CFU) were quantified on TSA10 supplemented with rifampicin. The averages ± sd of 3 replicates are plotted. Statistical analyses were performed using Dunn’s Multiple Comparison Test (* p-value < 0.05).

**Table S1. Strains and plasmids used in this study**

**Table S2. Primers used for T6SS mutant constructions**

