## Supplementary figures and images for "The Type VI secretion system *of Stenotrophomonas rhizophila* CFBP13503 limits the transmission of *Xanthomonas campestris* pv*. campestris* 8004 from radish seeds to seedlings"

### Figure S2

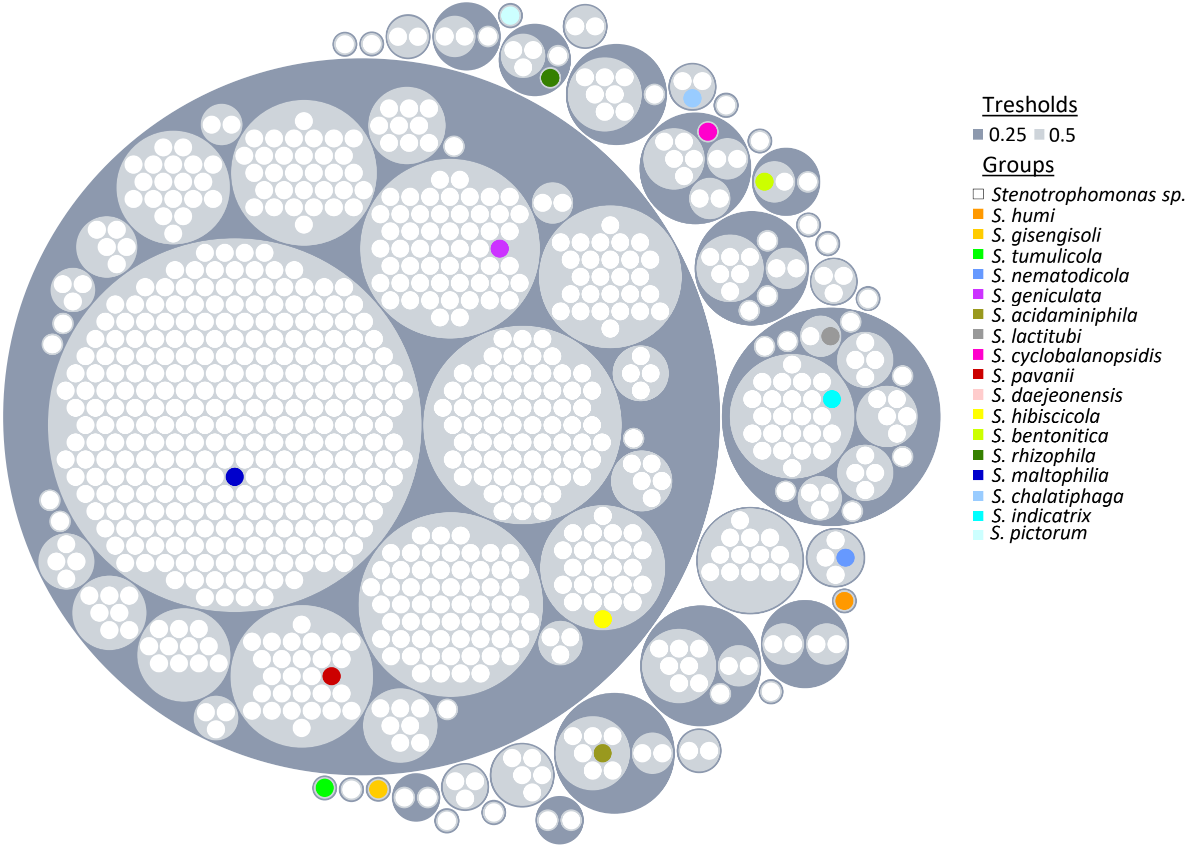

### Figure S3

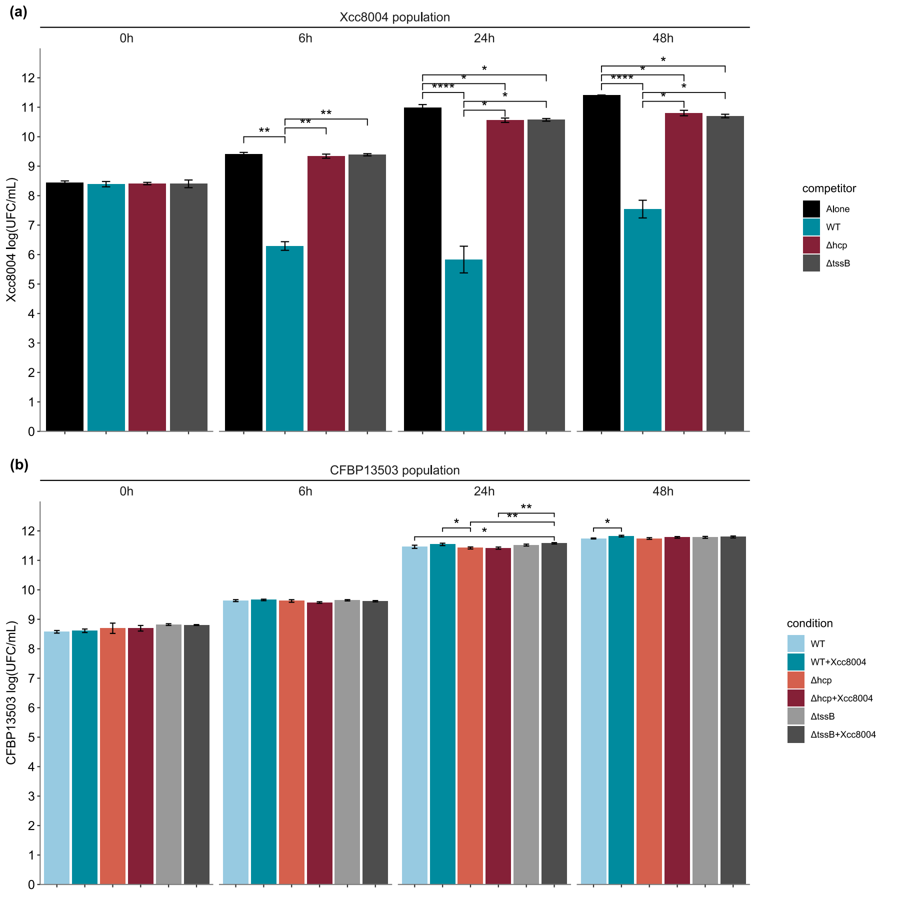

### Figure S4

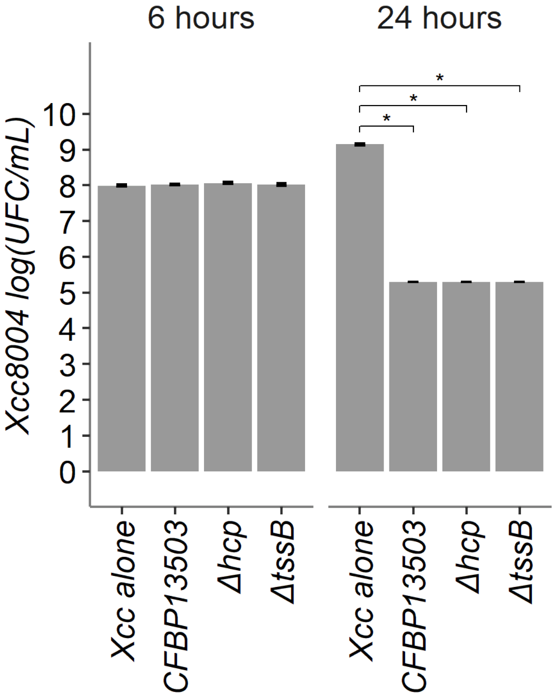
